# ibaqpy: A scalable Python package for baseline quantification in proteomics leveraging SDRF metadata

**DOI:** 10.1101/2025.02.08.637208

**Authors:** Ping Zheng, Enrique Audain, Henry Webel, Chengxin Dai, Joshua Klein, Marc-Phillip Hitz, Timo Sachsenberg, Mingze Bai, Yasset Perez-Riverol

## Abstract

Intensity-based absolute quantification (iBAQ) is essential in proteomics as it allows for the assessment of a protein’s absolute abundance in various samples or conditions. However, the computation of these values for increasingly large-scale and high-throughput experiments, such as those using DIA, TMT, or LFQ workflows, poses significant challenges in scalability and reproducibility. Here, we present ibaqpy (https://github.com/bigbio/ibaqpy), a Python package designed to compute iBAQ values efficiently for experiments of any scale. ibaqpy leverages the Sample and Data Relationship Format (SDRF) metadata standard to incorporate experimental metadata into the quantification workflow. This allows for automatic normalization and batch correction while accounting for key aspects of the experimental design, such as technical and biological replicates, fractionation strategies, and sample conditions. Designed for large-scale proteomics datasets, ibaqpy can also recompute iBAQ values for existing experiments when an SDRF is available. We showcased ibaqpy’s capabilities by reanalyzing 17 public proteomics datasets from ProteomeXchange, covering HeLa cell lines with 4,921 samples and 5,766 MS runs, quantifying a total of 11,014 proteins. In our reanalysis, ibaqpy is a key component in automating reproducible quantification, reducing manual effort and making quantitative proteomics more accessible while supporting FAIR principles for data reuse.

## 1. Introduction

Intensity-based absolute quantification (iBAQ) is a key technique in proteomics for estimating the absolute abundance of proteins in various samples, cell types, or biological conditions. This method standardizes protein quantification by normalizing the summed peptide signal intensities against the theoretical number of observable peptides, enabling consistent comparisons across experiments and conditions. In recent years, iBAQ has become instrumental in establishing baseline protein expression levels across diverse experiments, samples, and conditions. Baseline gene and protein expression refers to the measurement of RNA and protein levels in different tissues, cell types, and cell lines under normal or specific conditions [1-3]. The baseline expression, rather than being an absolute or unique value, can be understood as an expression profile—a pattern or range of expression within a specific tissue or cell line. These profiles offer valuable insights into the typical protein expression under specific conditions, such as within a particular tissue or cell type. Understanding baseline expression is crucial in drug development, as it provides insights into the availability and range of a target molecule’s expression in the tissue of interest [4], while systematic measurements of protein expression levels enhance our understanding of fundamental biological processes and inform the design of new therapeutic strategies [4, 5]. Resources like PaxDB (https://pax-db.org/) [5], ProteomicsDB (https://www.proteomicsdb.org/) [6] or quantms (https://quantms.org/home) [7] provide curated datasets that aggregate protein expression data across tissues and organisms, offering baseline protein expression profiles. While these resources may refer to expression profiles differently, they share the goal of providing a range of expression values of a particular protein on multiple conditions, tissues, disease or cell lines. For instance, PaxDB focuses on generating a consensus view of normal or healthy proteomes and represents protein abundance as ‘parts per million’ (ppm), normalised against all other protein molecules in the sample [5]. While ProteomicsDB and quantms use the normalized intensity-based absolute quantification values (iBAQ) of each protein on each specific condition.

iBAQ has proven highly reproducible in measuring protein abundance across experiments, even from different analytical methods [8-11], and for large dataset integration [2, 6, 12]. While tools like MaxQuant/MaxDIA [13, 14] and FragPipe [15] enable iBAQ computation for both multiplex and label-free experiments, most tools – especially those for DIA - like DIA-NN [16] or Spectronaut do not produce iBAQ values, but other metrics like MaxLFQ [17]. Although calculating the iBAQ value is straightforward, the process of transforming peptide feature intensities— a feature intensity is defined by the intensity of a peptidoform (modified sequence and its charge state) and retention time in a given MS run—into protein quantification values, along with the normalization steps, varies across tools and algorithms, impacting the final iBAQ results [18, 19]. In 2014, Rosenberger *et al*. [19] published the first package in R (aLFQ) to compute absolute quantification for multiple workflows, including OpenMS/OpenSWATH [20, 21] and Skyline [22] and multiple absolute methods such as iBAQ, APEX, and NSAF, providing a comprehensive framework for absolute label-free quantification. However, aLFQ does not scale well for large-scale experiments involving thousands of samples, as commonly seen in modern deep proteomics studies. Furthermore, it lacks integration with experimental design, such as sample information, including biological and technical replicates. Recently, the HUPO-PSI group released a new metadata format (SDRF) that captures the sample metadata but also the experimental design, including sample fractions and technical and biological replicates [23]. The SDRF has been widely adopted in multiple workflows, including quantms [2] and FragPipe [15].

Here, we present ibaqpy (https://github.com/bigbio/ibaqpy), a Python-based package to calculate iBAQ values and baseline expression profiles for large-scale proteomics experiments guided by metadata and the experimental design stored in SDRF files. By employing the SDRF annotated experimental design, which includes fractions, as well as technical and biological replicates, ibaqpy enables researchers to conduct proper normalisation at the peptide feature, peptide, and protein quantification steps. This metadata-driven approach ensures reproducible results while accounting for batch effects and other confounding variables, making SDRF integration a cornerstone for baseline iBAQ-based calculation across experiments. ibaqpy is designed for large-scale proteomics experiments where millions of features are quantified across thousands of samples. Additionally, various normalisation and sample processing methods are employed to convert features into protein quantification values via a metadata-informed workflow. We reanalyzed 17 large-scale HeLa datasets from ProteomeXchange [24], creating the largest corpus of HeLa protein expression profiles to support batch effect correction, protein co-expression studies, and deep learning algorithm development. A total of 5,766 MS runs were analyzed, and 11,014 proteins were quantified using ibaqpy and more than 200 million peptide features. ibaqpy is an open-source tool currently integrated into the quantms workflow [2, 9], providing a resource for computing iBAQ normalised values across all reanalyses of baseline datasets for multiple analytical methods, including TMT/ DDA-LFQ and DIA.

## 2. Methods

### 2.1. Hela datasets

A total of 17 publicly available HeLa proteomic datasets from the ProteomeXchange [24] and PRIDE Archive [25] were selected to benchmark and develop ibaqpy; and construct a comprehensive corpus of Hela baseline protein expression (**Table 1**). All datasets were annotated using the SDRF [23] file format, and the final results were converted to quantms.io format (https://github.com/bigbio/quantms.io), a Parquet-based format that enables sharing the peptide features, protein intensities, and absolute expression values for each experiment. The dataset PXD042233 is the largest reanalyzed [26]. In this case, instead of analysing the original 7,410 deposited, a subset of only 4,359 samples were analyzed (**Supplementary Note 1**).

**Table 1:** Overview of datasets analyzed using quantms and their corresponding reanalyses. Certain datasets underwent multiple reanalyses due to variations in experimental designs, such as DIA versus DDA, or differences in sample preparation, including using distinct enzymes. Some datasets do not have ProteomeXchange accession; in those cases, the PubMed accession or the accession of the original database will be used. The total number of quantified proteins refers to the proteins measured in at least one sample across all experiments.

| Accession | Reanalysis Accession | # Peptide Features | # Quantified Proteins | # Samples | # MS run | Analytical method |
| --- | --- | --- | --- | --- | --- | --- |
| PMID22068331 | PMID22068331.1 | 573'912 | 6096 | 10 | 70 | DDA-LFQ |
| PMID22068331 | PMID22068331.2 | 580'839 | 5262 | 12 | 74 | DDA-LFQ |
| PMID22068331 | PMID22068331.3 | 386'689 | 5764 | 8 | 48 | DDA-LFQ |
| PXD000269 | PXD000269 | 959'155 | 7903 | 4 | 24 | DDA-LFQ |
| PXD000396 | PXD000396 | 632'587 | 3096 | 12 | 40 | DDA-FLQ |
| PXD001305 | PXD001305 | 2'314'837 | 3847 | 55 | 102 | DDA-LFQ |
| PXD004452 | PXD004452 | 1'122'748 | 8519 | 6 | 230 | DDA-LFQ |
| PXD004940 | PXD004940 | 1'294'806 | 5434 | 15 | 45 | DDA-LFQ |
| PXD005481 | PXD005481 | 688'759 | 2566 | 17 | 51 | DDA-LFQ |
| PXD009273 | PXD009273.1 | 5'330'815 | 4344 | 141 | 141 | DIA-LFQ |
| PXD009875 | PXD009875 | 1'205'335 | 4583 | 17 | 104 | DDA-LFQ |
| PXD010150 | PXD010150 | 581'760 | 5709 | 10 | 46 | DDA-LFQ |
| PXD013658 | PXD013658.1 | 2'920'665 | 9037 | 4 | 12 | DIA-LFQ |
| PXD013658 | PXD013658.2 | 517'069 | 9540 | 1 | 20 | DDA-LFQ |
| PXD014058 | PXD014058 | 793'309 | 8447 | 4 | 80 | DDA-LFQ |
| PXD030406 | PXD030406 | 2'086'702 | 4595 | 16 | 48 | DDA-LFQ |
| PXD039414 | PXD039414 | 305'328 | 1975 | 17 | 17 | DIA-LFQ |
| PXD042233 | PXD042233 | 155'660'528 | 4452 | 4359 | 4359 | DDA-LFQ |
| PXD054015 | PXD054015.1 | 6'341'144 | 7864 | 21 | 63 | DIA-LFQ |
| PXD048325 | PXD048325 | 25'683'725 | 7717 | 192 | 192 | DIA-LFQ |
| **# datasets: 17, # reanalyses 20** |  | **209'980'712** | **11014\*** | **4921** | **5766** |  |

### 2.2. Data analysis and feature maps

quantms (https://github.com/bigbio/quantms/) is an open-source and cloud-based nextflow workflow that enables the reanalysis of public proteomics data [2]. The version of quantms used in the present study was 1.3.1dev, which uses the newest version of the DIA-NN tool (1.9.2) [16], and for the LFQ-DDA analysis uses multiple search engines COMET, MSGF+ and SAGE supported by the latest development version of OpenMS (version 3.3) [20].

### 2.3 Batch effect correction

Batch effect correction was performed using the Python library *inmoose* (https://github.com/epigenelabs/inmoose) [27, 28]. The current implementation (v0.7.3) supports two methods for batch correction: *combat-norm* and *combat-seq. combat-norm* [29] assumes a normal distribution for the underlying data, which is closer to MS intensity distributions, while *combat-seq* is suitable for count-based data like RNA-seq, assuming a negative binomial distribution of the data [27]. As therefore *combat-seq* cannot be used with MS-based intensity data, we only used and integrated *combat-norm* in ibaqpy. We integrate *combat-norm* (*inmoose*) in ibaqpy, and in the present analysis, we used the default parameters.

## 3. Results and Discussion

### 3.1. ibaqpy workflow: from peptide features to iBAQ values

**Figure 1a** illustrates the main steps performed by the ibaqpy package: 1) feature preprocessing, 2) normalization of fractions and technical replicates, 3) peptide level normalization, 4) feature-to-peptide normalization, and 5) the peptides2protein quantification step (**Figure 1a**). The library requires three key files: a Parquet file containing all quantified peptide features in the quantms.io format (see https://github.com/bigbio/quantms.io/blob/main/docs/README.adoc) (**Figure 1b-c**); an SDRF file specifying the experimental design; and a FASTA file with the protein sequences used for quantification. The feature preprocessing step enables users to exclude decoys, contaminants, entrapments, or proteins that are significantly overexpressed in the sample, which could bias downstream analyses such as normalization—for example, plasma samples contaminated with other blood cell types [30, 31]. Additionally, users can filter out proteins supported by fewer than a specified number of peptides or unique peptides.

**Figure 1:**
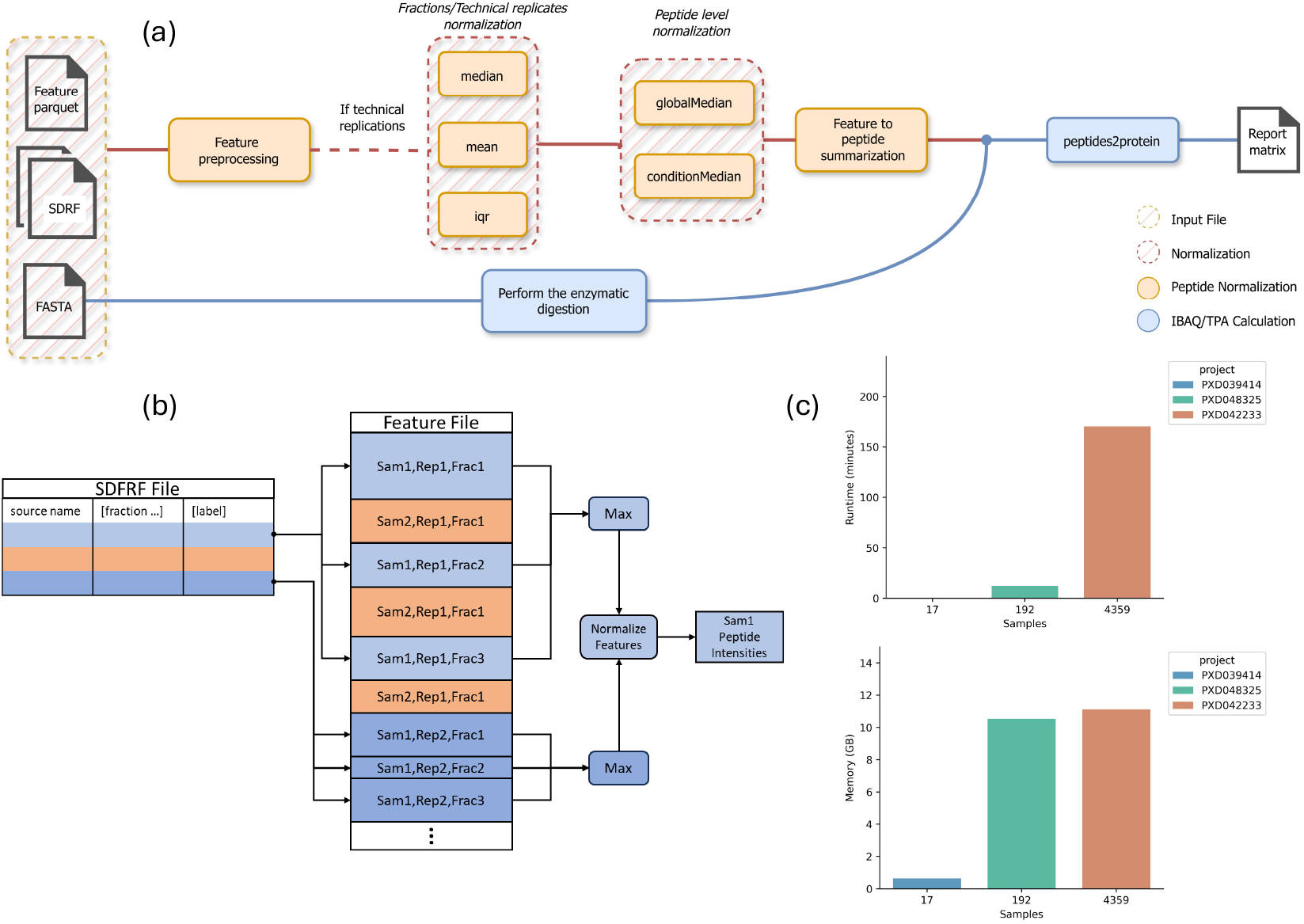
(**a)** The ibaqpy workflow starts with a feature Parquet file, an SDRF sample metadata file, and a FASTA protein database. The first preprocessing step removes features belonging to multiple proteins like contaminants, decoys or entrapments. In the preprocessing component, peptide features undergo multiple preprocessing steps, such as median, mean, or interquartile range (IQR) calculations, before being summarized using global or condition-specific medians. This is followed by peptide normalization and the computation of the baseline/absolute values such as iBAQ or normalized iBAQ (riBAQ) or TPA. The quantms feature file contains a table of peptidoform ion feature intensities measured for each MS experiment, and each experiment is associated with a source sample, replication level, fraction, and/or multiplexing channels (if used) defined in an SDRF file. The experimental design metadata is then used to decide how to aggregate peptidoform features for normalization. **(c)** Performance evaluation, memory (GB) and runtime (minutes) of ibaqpy across three different projects with varying sample sizes: PXD039414 (17 samples), PXD048325 (192 samples), and PXD042233 (4359 samples).

Peptide features are first normalized across fractions and technical replicates using one of three methods: median, mean, or interquartile range (IQR). By default, the tool uses the median method. Figure 2a illustrates the impact of median normalization on feature intensities when applied to dataset PXD030406 (**Supplementary Note 2**). A second normalization step is needed to transform peptide features – all peptidoforms for a given peptide, including modifications and charge states - into peptide intensities. At a peptide level, two normalization methods can be applied, (i) the global median: calculating the median for each sample and then taking the median of these medians as the global median to correct the distribution of all samples; (ii) the condition median: calculating the median for each sample under the same condition and then taking the average of these medians as the condition median to correct the distribution of all samples under that condition. Both methods use the SDRF information to capture the condition (factor value in the SDRF) and the corresponding fraction, technical and biological replicates information (**Figure 1a**).

**Figure 2:**
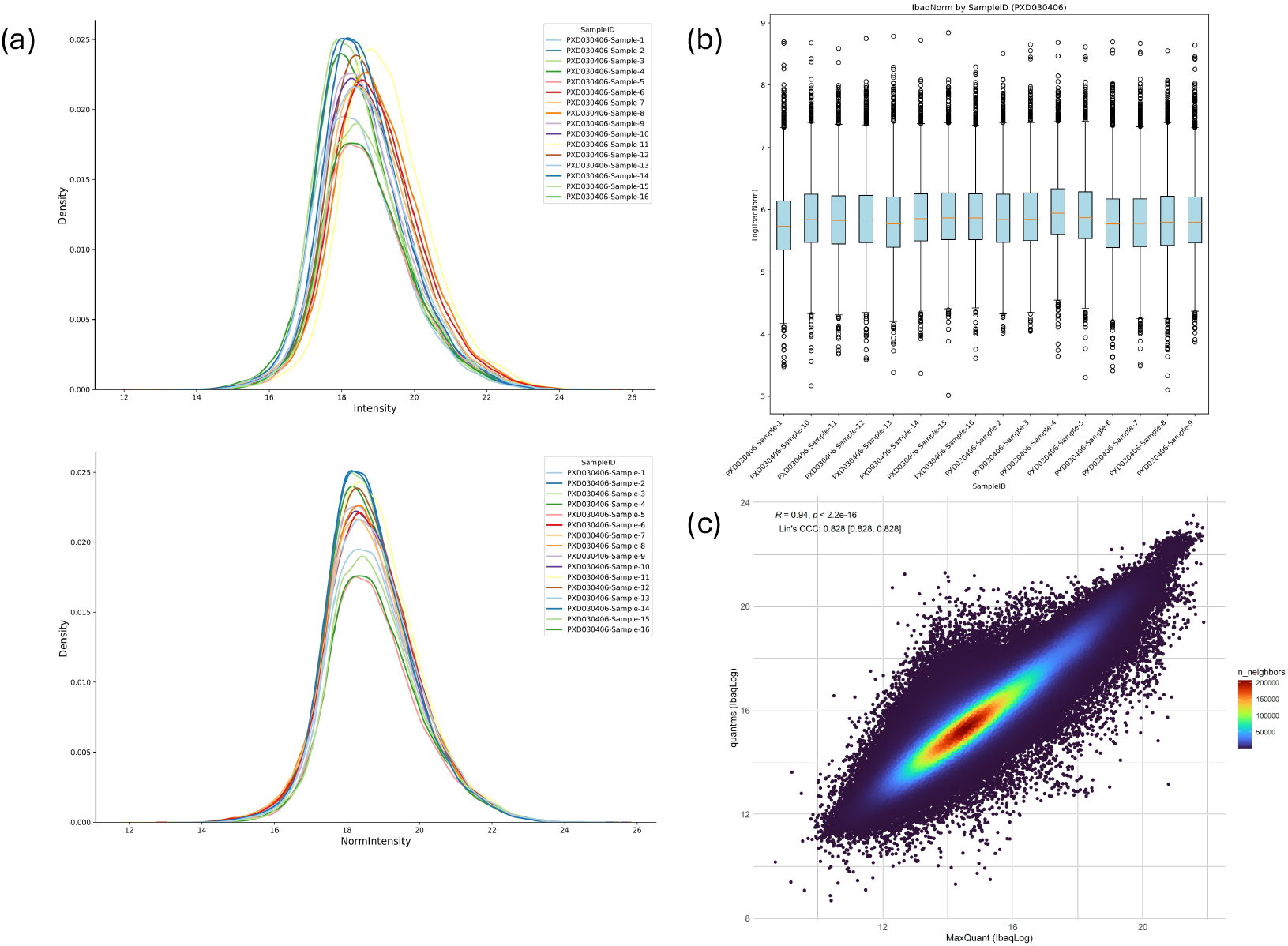
**(a)** Density plots illustrate the distribution of peptide feature intensities before (left panel) and after (right panel) normalization using ibaqpy for project PXD030406. **(b)** Box plots visualize the variability of protein IbaqLog for all samples in project PXD030406 within each group. **(c)** Correlation between log iBAQ values computed with ibaqpy for 500 samples of dataset PXD042233, starting from peptide features quantified with MaxQuant (as originally submitted to PRIDE) and quantms (after reanalysis), with the Pearson correlation coefficient (R) and Lin’s concordance correlation coefficient (CCC) quantifying the agreement.

The iBAQ value is calculated by dividing the total intensity of a protein’s peptides by the theoretical number of peptides generated from experimental protease treatment, i.e. trypsin digestion, accounting for protein size and enabling abundance comparisons within a sample [8, 9]. Theoretical peptides are typically 6–30 amino acids, excluding missed cleavages and modifications. Apart from computing the iBAQ value for each protein (Ibaq); ibaqpy computes three normalized iBAQ metrics:

- IbaqNorm (riBAQ): iBAQ divided by the total iBAQ across all proteins in the sample [32].
- IbaqLog: Log10 transformation of IbaqNorm scaled by a constant of 10, used in ProteomicsDB [6] and quantms [2].
- IbaqPpb: IbaqNorm scaled to parts-per-billion (PPB) for easier interpretation [3]. Additionally, ibaqpy calculates absolute expression values like:
- Total Protein Approach (TPA): calculates protein abundance by dividing the summed peptide intensities of a protein by the total intensity of all detected proteins in the sample [33].
- Cellular Protein Copy Number: estimating protein copies per cell the protein abundance, molecular mass, and Avogadro’s number [34].

Figure 2b shows the distribution of IbaqLog values across all 16 samples from dataset PXD030406. After normalizing feature intensities (Figure 2a) and scaling iBAQ values by the total iBAQ within each sample (IbaqLog), this variability is notably reduced, confirming observations previously made by our team [3, 9]. All metrics computed by ibaqpy, following the normalisation of feature and peptide intensities, exhibit lower variability across samples, similar to IbagLog (**Supplementary Figure 4** – Copy cell number for PXD030406).

#### 3.1.1. Parquet-based feature files for large-scale experiments

The peptide feature format is an Apache Parquet-based file (https://github.com/bigbio/quantms.io/blob/main/docs/README.adoc) that contains the peptide sequence, modifications, charge state, intensity, the MS run where the feature has been identified, and the sample accession from the SDRF (**Figure 1b**). The feature Parquet file allows for the efficient storage of millions of features across thousands of samples, ensuring scalability for large datasets (**Figure 1b**). During the feature normalization step, the process is optimized to scale for experiments with a large number of samples while maintaining consistent performance without adversely affecting CPU or memory usage (**Figure 1c**).

DuckDB (https://duckdb.org/) provides native support for the Parquet file format. DuckDB and SQL-based language are used to query the feature file for the fields that are needed by the tool: peptidoform, charge, intensity, MS run, label, and sample accession. DuckDB aims to fully utilize available memory to accelerate query processing while managing datasets that exceed memory (**Supplementary Note 4**). ibaqpy utilizes DuckDB to retrieve data using samples as indices and the median calculation for normalization. ibaqpy loads only a fixed number of samples into memory at a time, ensuring that it can perform data processing operations even in environments with limited memory.

#### 3.1.2. SDRF-based quantification

Leveraging the Sample and Data Relationship Format (SDRF) [23] to compute iBAQ values with ibaqpy introduces a metadata-driven approach to protein quantification that offers significant advantages over traditional methods (**Figure 1b**). SDRF provides machine-readable, structured metadata that includes fractions, technical replicates, and biological replicates, enabling ibaqpy to systematically incorporate experiment structure into the quantification workflow. Unlike other software like aLFQ [19], which computes iBAQ values independently for each MS run, ibaqpy can process multiple runs in the context of the experimental design. Another key advantage of SDRF-based quantification is its alignment with FAIR principles [35], promoting data interoperability and reusability of public proteomics data.

Researchers can convert existing public datasets in PRIDE Archive to peptide features files using the quantms.io toolkit, remap the proteins to a new version of the UniProt database and recompute iBAQ values using ibaqpy after annotating the SDRF file (read the documentation about how to convert PRIDE Archive submissions to quantms.io - https://github.com/bigbio/quantms.io/blob/main/docs/ibaq_usecase.adoc). Figure 2c illustrates the correlation of IbaqLog values for 500 samples from the dataset PXD042233. The iBAQ values were computed using the original peptide features obtained with MaxQuant (as submitted to PRIDE Archive) and those derived from reanalysis with quantms. The Pearson correlation between the two analyses, calculated using ibaqpy, is 0.94, alongside a high Lin’s concordance correlation coefficient of 0.82. Consistent with our previous findings [9] —where we compared iBAQ values across TMT and LFQ quantification methods for the same samples, as well as iBAQ values from ProteomicsDB and quantms—we demonstrate that iBAQ values calculated with ibaqpy using identical normalization methods remain highly correlated, even when different workflows are used for peptide quantification.

### 3.2. A corpus of Hela expression profiles

We re-annotated 17 large-scale Hela public datasets. To standardize the annotations, we use the SDRF format, the SDRF Cell Line Metadata Database (https://github.com/bigbio/sdrf-cellline-metadata-db), which enables annotation for every sample in the same way for all the properties of the cell line, including sampling site, organism part, and Cellosaurus accession [35]. A total of 5,766 MS runs were analyzed from 4,921 samples, and 20 reanalyses were performed (**Table 1**). All reanalyses were performed with quantms, 5% PSM FDR and 1% FDR at the protein level; for each protein, at least two unique peptides were needed. In total, we quantified 209,980,712 peptide features and 11,014 proteins across the entire collection.

**Figure 3a** illustrates the distribution of proteins quantified across the 17 projects. Notably, 11% of the identified proteins (n = 1,224) are only quantified in a single project, whereas the majority, comprising 89%, are quantified in multiple projects. Furthermore, it is significant to highlight that merely 6% of the proteins are quantified in only a single sample (**Figure 2b**). In contrast, a substantial 94% of the proteins are both identified and quantified in more than one sample, making this one of the most reproducible and high-quality collections of quantified proteins, as nearly every protein quantified with at least two unique peptides is consistently observed across multiple experiments and samples. In addition to quantifying proteins across numerous experiments and samples, **Figure 3c** illustrates the strong correlation of iBAQ values for each protein across all experimental conditions. The lower correlation between the two projects is 0.56, while the majority of projects show a correlation higher than 0.89 (Figure 3c). PXD014058 shows a lower correlation compared to all the other projects reanalyzed. Finally, as anticipated, **Figure 3d** illustrates that as the number of samples increases, the median iBAQ may also relate to the observation that the most abundant proteins are easily detected and quantified alongside the less abundant proteins.

**Figure 3:**
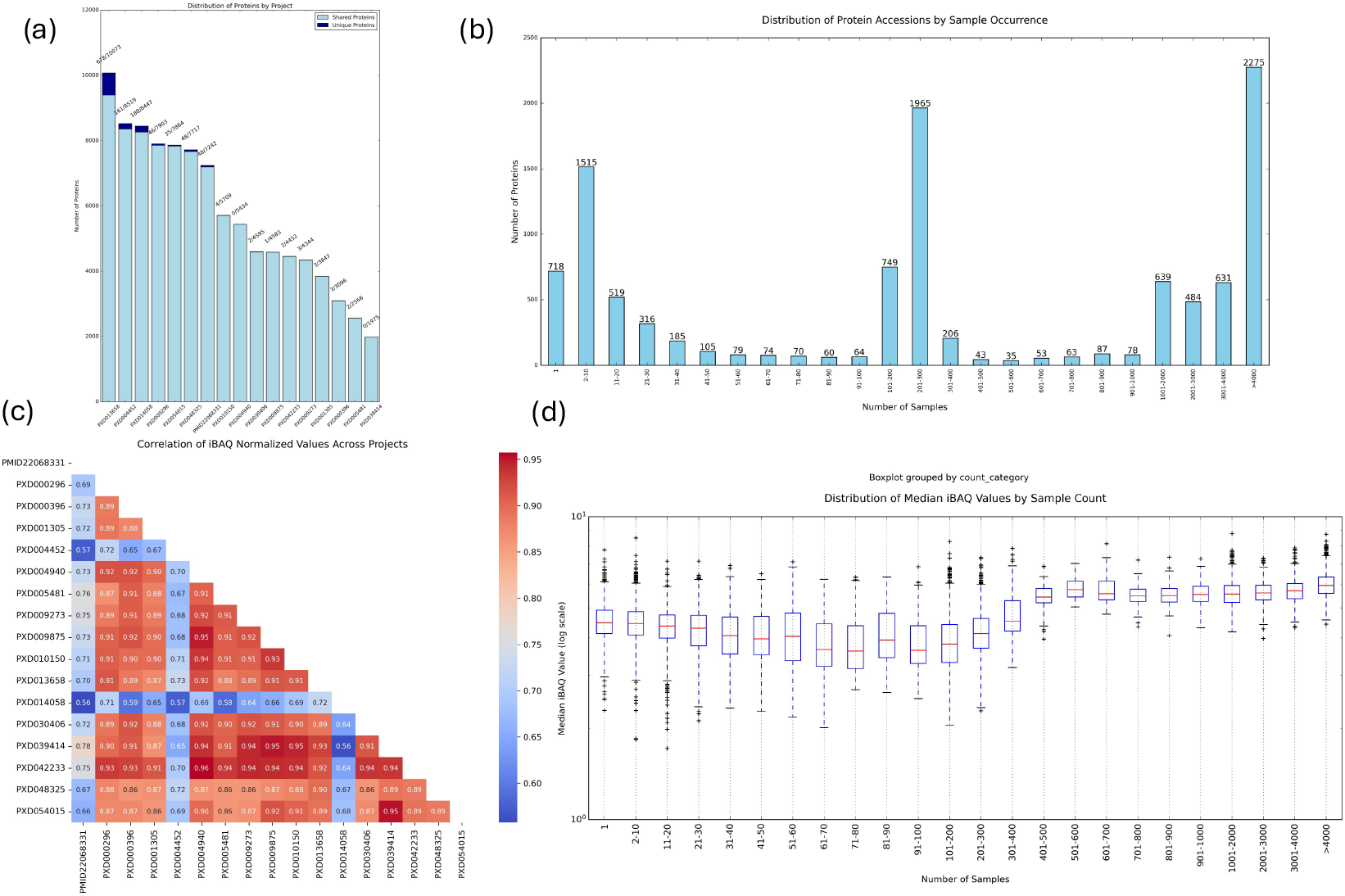
**(a)** Bar plot showing the number of total and unique proteins quantified in each of the 17 projects. **(b)** Bar plot showing the distribution of proteins quantified in different numbers of samples. **(c)** Heatmap showing the correlation of the normalized iBAQ for the 17 projects reanalyzed. **(d)** Boxplot showing the distribution of median iBAQ values for proteins quantified in different numbers of samples.

#### 3.2.1 Large-scale batch effect correction on baseline data

After data aggregation, intra- and inter-experimental variability is expected due to differences in experimental conditions, as observed in **Figure 4a**. This variability can impact expression profiles within and across experiments, potentially confounding downstream analyses. To address this, we applied the *combat-norm* batch effect correction method, implemented in the *inmoose* library [27, 28] and is now part of the ibaqpy workflow. Similar to other intensity-based quantification techniques like RNA microarrays [29], iBAQ values are susceptible to batch effects from sample processing variability. We chose unnormalized iBAQ values for batch effect correction using combat-norm to prevent distortions introduced by previous normalization steps (e.g. riBAQ). Batch effect correction significantly reduces variability across samples (**Figure 4b**) and enhances consistency across experiments. The correction was less effective in those projects with the highest number of unique proteins (**Figure 3a**), projects PXD013658 and PXD004452. The effectiveness of batch effect correction was further evaluated using the Silhouette score and Entropy of batch mixing (EBM) (**Supplementary Note 5)**. Both metrics indicated improved mixing of samples across batches, supporting the robustness of the correction approach. The observed results align with previous proteomics studies that utilized these metrics to evaluate the batch effect correction process [36, 37]. To assess the impact of the batch correction on specific protein expression profiles, we compared the expression of low- and high-abundance proteins before and after correction (**Figures 4c** and **4d)**. The batch correction procedure harmonised the expression profiles while preserving the underlying data structure.

**Figure 4.**
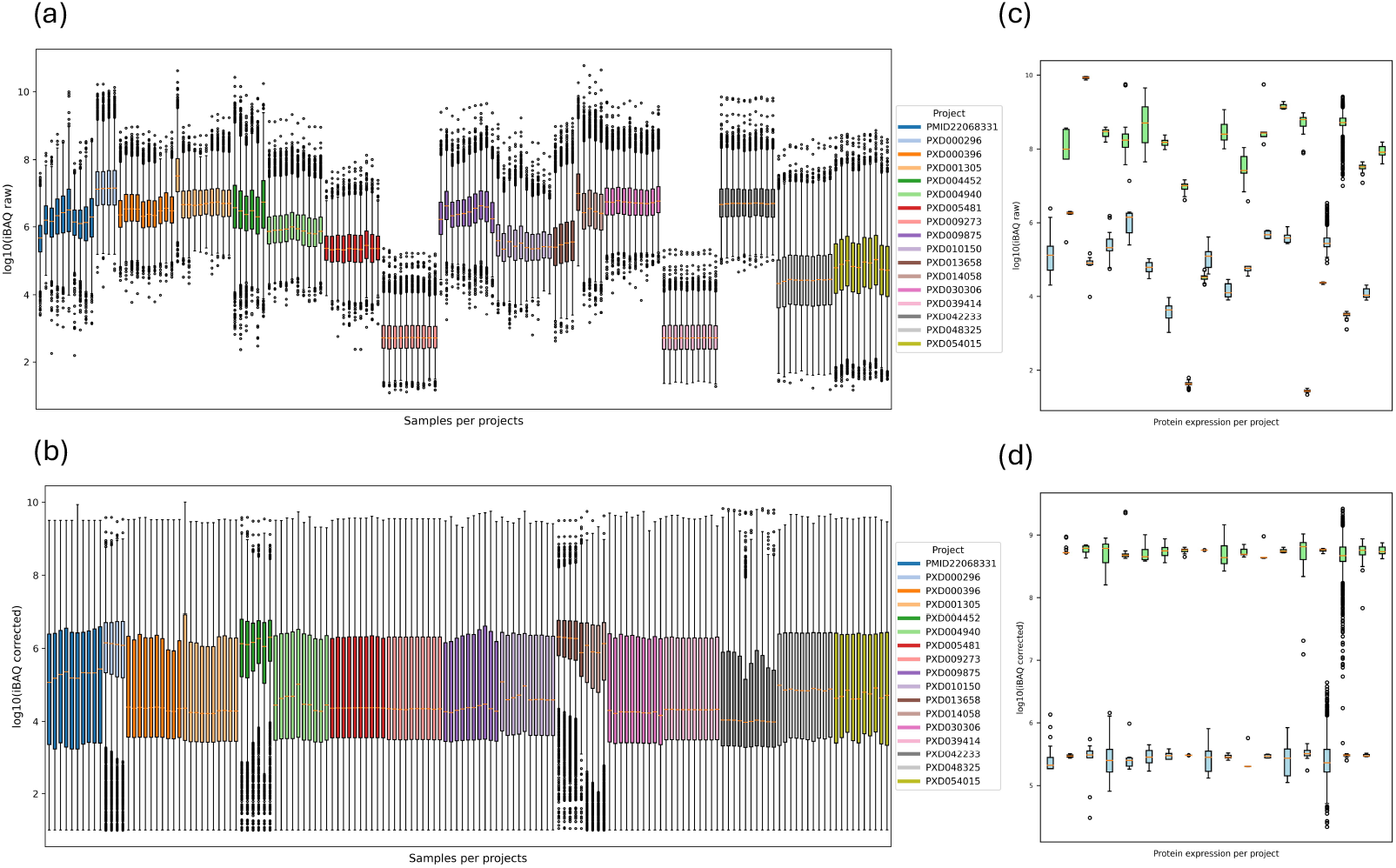
Evaluation of the batch effect correction in the complete dataset. **(a)** Raw iBAQ distribution per sample. **(b)** Batch-corrected iBAQ distribution per sample. Boxplots display a maximum of 10 samples per project. Per project raw **(c)** and corrected **(d)** iBAQ distributions of two proteins expressed in the lowest (Q5T4S7 in blue) and highest (P05387 in green) quartile. The y-axis is shown in log-transformed scales.

### 3.3 Data and Code Availability

All the reanalyses performed in this study, including the SDRFs annotated, the peptide-spectrum-matches, and outputs from quantms can be searched and retrieved from quantms.org. Additionally, the quantms.io representation of every reanalysis can be found in the following ftp (https://ftp.pride.ebi.ac.uk/pub/databases/pride/resources/proteomes/ibaqpy-research/). Ibaqpy is an open-source Python library (https://github.com/bigbio/ibaqpy) and the specification of quantms.io can be found on GitHub (https://github.com/bigbio/quantms.io).

## 4. Conclusion

iBAQ absolute quantification and baseline expression profiles are becoming fundamental for understanding biological systems and advancing drug development. ibaqpy represents a significant step forward in enabling scalable, metadata-driven computation of iBAQ values for large-scale proteomics experiments. By leveraging Parquet-based feature files and SDRF metadata, ibaqpy not only provides robust, experimental design-aware normalization and quantification methods but also ensures reproducibility following FAIR principles. ibaqpy’s integration into the quantms workflow further solidifies its utility in large-scale reanalyses and highlights its alignment with community standards for open and reproducible research. Moving forward, ibaqpy has the potential to become a standard tool for absolute quantification in proteomics, enabling the generation of high-quality baseline expression maps that benefit both fundamental and applied biological research. These maps will serve as valuable resources for a wide range of studies.

Our reanalysis of 17 HeLa datasets, comprising 5,766 MS runs and quantifying over 11,000 proteins, showcases the scalability and performance of ibaqpy, generating one of the most comprehensive protein expression corpora available. ibaqpy benefits from SDRF’s compatibility with public proteomics repositories like PRIDE, facilitating seamless integration into submission workflows and downstream analyses. The use of SDRF automates the parsing of metadata, reducing manual effort and user error while enabling better cross-study comparisons. Beyond its current capabilities, ibaqpy has the potential to evolve into a comprehensive framework for metadata-informed data processing. This could support advanced normalization strategies, sophisticated batch correction techniques, and other improvements, further enhancing its role in large-scale proteomics research.

## Acknowledgements

We want to thank Julianus Pfeuffer, developer of quantms and the first iteration of ibaqpy. Ping Zheng and Mingze Bai are supported by Chongqing’s Science and Technology Innovation Key R&D Program (CSTB2023TIAD-STX0002). Yasset Perez-Riverol would like to acknowledge funding from EMBL core funding, Wellcome grants (208391/Z/17/Z, 223745/Z/21/Z) and the BBSRC grant ‘DIA-Exchange’ [BB/X001911/1]. T.S. acknowledges funding by the Federal Ministry of Education and Research in the frame of de.NBI/ELIXIR-DE (W-de.NBI-022) and is supported by the Ministry of Science, Research and Arts Baden-Württemberg.

## References

[1] R. Petryszak, T. Burdett, B. Fiorelli, N.A. Fonseca, M. Gonzalez-Porta, E. Hastings, W. Huber, S. Jupp, M. Keays, N. Kryvych, J. McMurry, J.C. Marioni, J. Malone, K. Megy, G. Rustici, A.Y. Tang, J. Taubert, E. Williams, O. Mannion, H.E. Parkinson, A. Brazma, Expression Atlas update--a database of gene and transcript expression from microarray- and sequencing-based functional genomics experiments, Nucleic Acids Res 42(Database issue) (2014) D926–32.

[2] C. Dai, J. Pfeuffer, H. Wang, P. Zheng, L. Käll, T. Sachsenberg, V. Demichev, M. Bai, O. Kohlbacher, Y. Perez-Riverol, quantms: a cloud-based pipeline for quantitative proteomics enables the reanalysis of public proteomics data, Nature Methods (2024).

[3] A.F. Jarnuczak, H. Najgebauer, M. Barzine, D.J. Kundu, F. Ghavidel, Y. Perez-Riverol, I. Papatheodorou, A. Brazma, J.A. Vizcaino, An integrated landscape of protein expression in human cancer, Sci Data 8(1) (2021) 115.

[4] E.M. McDonagh, G. Trynka, M. McCarthy, E.R. Holzinger, S. Khader, N. Nakic, X. Hu, H. Cornu, I. Dunham, D. Hulcoop, Human Genetics and Genomics for Drug Target Identification and Prioritization: Open Targets’ Perspective, Annu Rev Biomed Data Sci 7(1) (2024) 59–81.

[5] Q. Huang, D. Szklarczyk, M. Wang, M. Simonovic, C. von Mering, PaxDb 5.0: Curated Protein Quantification Data Suggests Adaptive Proteome Changes in Yeasts, Mol Cell Proteomics 22(10) (2023) 100640.

[6] L. Lautenbacher, P. Samaras, J. Muller, A. Grafberger, M. Shraideh, J. Rank, S.T. Fuchs, T.K. Schmidt, M. The, C. Dallago, H. Wittges, B. Rost, H. Krcmar, B. Kuster, M. Wilhelm, ProteomicsDB: toward a FAIR open-source resource for life-science research, Nucleic Acids Res 50(D1) (2022) D1541–D1552.

[7] C. Dai, J. Pfeuffer, H. Wang, P. Zheng, L. Kall, T. Sachsenberg, V. Demichev, M. Bai, O. Kohlbacher, Y. Perez-Riverol, quantms: a cloud-based pipeline for quantitative proteomics enables the reanalysis of public proteomics data, Nat Methods 21(9) (2024) 1603–1607.

[8] B. Schwanhausser, D. Busse, N. Li, G. Dittmar, J. Schuchhardt, J. Wolf, W. Chen, M. Selbach, Global quantification of mammalian gene expression control, Nature 473(7347) (2011) 337–42.

[9] H. Wang, C. Dai, J. Pfeuffer, T. Sachsenberg, A. Sanchez, M. Bai, Y. Perez-Riverol, Tissue-based absolute quantification using large-scale TMT and LFQ experiments, Proteomics 23(20) (2023) e2300188.

[10] J. Wang, W. Yu, R. D’Anna, A. Przybyla, M. Wilson, M. Sung, J. Bullen, E. Hurt, G. D’Angelo, B. Sidders, Z. Lai, W. Zhong, Pan-Cancer Proteomics Analysis to Identify Tumor-Enriched and Highly Expressed Cell Surface Antigens as Potential Targets for Cancer Therapeutics, Mol Cell Proteomics 22(9) (2023) 100626.

[11] J. Wang, X. Tian, W. Yu, B. Pullman, J. Bullen Jr, E. Hurt, W. Zhong, Evaluating computational approaches for comparison of protein expression across cancer indications, bioRxiv (2024) 2024.08. 26.609731.

[12] C. Schaab, T. Geiger, G. Stoehr, J. Cox, M. Mann, Analysis of high accuracy, quantitative proteomics data in the MaxQB database, Mol Cell Proteomics 11(3) (2012) M111 014068.

[13] J. Cox, M. Mann, MaxQuant enables high peptide identification rates, individualized p.p.b.-range mass accuracies and proteome-wide protein quantification, Nat Biotechnol 26(12) (2008) 1367–72.

[14] P. Sinitcyn, H. Hamzeiy, F. Salinas Soto, D. Itzhak, F. McCarthy, C. Wichmann, M. Steger, U. Ohmayer, U. Distler, S. Kaspar-Schoenefeld, N. Prianichnikov, S. Yilmaz, J.D. Rudolph, S. Tenzer, Y. Perez-Riverol, N. Nagaraj, S.J. Humphrey, J. Cox, MaxDIA enables library-based and library-free data-independent acquisition proteomics, Nat Biotechnol 39(12) (2021) 1563–1573.

[15] A.T. Kong, F.V. Leprevost, D.M. Avtonomov, D. Mellacheruvu, A.I. Nesvizhskii, MSFragger: ultrafast and comprehensive peptide identification in mass spectrometry-based proteomics, Nat Methods 14(5) (2017) 513–520.

[16] V. Demichev, C.B. Messner, S.I. Vernardis, K.S. Lilley, M. Ralser, DIA-NN: neural networks and interference correction enable deep proteome coverage in high throughput, Nat Methods 17(1) (2020) 41–44.

[17] J. Cox, M.Y. Hein, C.A. Luber, I. Paron, N. Nagaraj, M. Mann, Accurate proteome-wide label-free quantification by delayed normalization and maximal peptide ratio extraction, termed MaxLFQ, Mol Cell Proteomics 13(9) (2014) 2513–26.

[18] M. Shi, C.A. Evans, J.L. McQuillan, J. Noirel, J. Pandhal, LFQRatio: A Normalization Method to Decipher Quantitative Proteome Changes in Microbial Coculture Systems, J Proteome Res 23(3) (2024) 999–1013.

[19] G. Rosenberger, C. Ludwig, H.L. Rost, R. Aebersold, L. Malmstrom, aLFQ: an R-package for estimating absolute protein quantities from label-free LC-MS/MS proteomics data, Bioinformatics 30(17) (2014) 2511–3.

[20] J. Pfeuffer, C. Bielow, S. Wein, K. Jeong, E. Netz, A. Walter, O. Alka, L. Nilse, P.D. Colaianni, D. McCloskey, J. Kim, G. Rosenberger, L. Bichmann, M. Walzer, J. Veit, B. Boudaud, M. Bernt, N. Patikas, M. Pilz, M.P. Startek, S. Kutuzova, L. Heumos, J. Charkow, J.C. Sing, A. Feroz, A. Siraj, H. Weisser, T.M.H. Dijkstra, Y. Perez-Riverol, H. Rost, O. Kohlbacher, T. Sachsenberg, OpenMS 3 enables reproducible analysis of large-scale mass spectrometry data, Nat Methods 21(3) (2024) 365–367.

[21] H.L. Rost, G. Rosenberger, P. Navarro, L. Gillet, S.M. Miladinovic, O.T. Schubert, W. Wolski, B.C. Collins, J. Malmstrom, L. Malmstrom, R. Aebersold, OpenSWATH enables automated, targeted analysis of data-independent acquisition MS data, Nat Biotechnol 32(3) (2014) 219–23.

[22] L.K. Pino, B.C. Searle, J.G. Bollinger, B. Nunn, B. MacLean, M.J. MacCoss, The Skyline ecosystem: Informatics for quantitative mass spectrometry proteomics, Mass Spectrom Rev 39(3) (2020) 229–244.

[23] C. Dai, A. Fullgrabe, J. Pfeuffer, E.M. Solovyeva, J. Deng, P. Moreno, S. Kamatchinathan, D.J. Kundu, N. George, S. Fexova, B. Gruning, M.C. Foll, J. Griss, M. Vaudel, E. Audain, M. Locard-Paulet, M. Turewicz, M. Eisenacher, J. Uszkoreit, T. Van Den Bossche, V. Schwammle, H. Webel, S. Schulze, D. Bouyssie, S. Jayaram, V.K. Duggineni, P. Samaras, M. Wilhelm, M. Choi, M. Wang, O. Kohlbacher, A. Brazma, I. Papatheodorou, N. Bandeira, E.W. Deutsch, J.A. Vizcaino, M. Bai, T. Sachsenberg, L.I. Levitsky, Y. Perez-Riverol, A proteomics sample metadata representation for multiomics integration and big data analysis, Nat Commun 12(1) (2021) 5854.

[24] E.W. Deutsch, N. Bandeira, Y. Perez-Riverol, V. Sharma, J.J. Carver, L. Mendoza, D.J. Kundu, S. Wang, C. Bandla, S. Kamatchinathan, S. Hewapathirana, B.S. Pullman, J. Wertz, Z. Sun, S. Kawano, S. Okuda, Y. Watanabe, B. MacLean, M.J. MacCoss, Y. Zhu, Y. Ishihama, J.A. Vizcaino, The ProteomeXchange consortium at 10 years: 2023 update, Nucleic Acids Res 51(D1) (2023) D1539–D1548.

[25] Y. Perez-Riverol, C. Bandla, D.J. Kundu, S. Kamatchinathan, J. Bai, S. Hewapathirana, N.S. John, A. Prakash, M. Walzer, S. Wang, J.A. Vizcaino, The PRIDE database at 20 years: 2025 update, Nucleic Acids Res 53(D1) (2025) D543–D553.

[26] H. Webel, Y. Perez-Riverol, A.B. Nielsen, S. Rasmussen, Mass spectrometry-based proteomics data from thousands of HeLa control samples, Sci Data 11(1) (2024) 112.

[27] A. Behdenna, M. Colange, J. Haziza, A. Gema, G. Appe, C.A. Azencott, A. Nordor, pyComBat, a Python tool for batch effects correction in high-throughput molecular data using empirical Bayes methods, BMC Bioinformatics 24(1) (2023) 459.

[28] M. Colange, G. Appé, L. Meunier, S. Weill, A. Nordor, A. Behdenna, Differential Expression Analysis with InMoose, the Integrated Multi-Omic Open-Source Environment in Python, bioRxiv (2024) 2024.11.14.623578.

[29] J.T. Leek, R.B. Scharpf, H.C. Bravo, D. Simcha, B. Langmead, W.E. Johnson, D. Geman, K. Baggerly, R.A. Irizarry, Tackling the widespread and critical impact of batch effects in high-throughput data, Nat Rev Genet 11(10) (2010) 733–9.

[30] I. Metatla, K. Roger, C. Chhuon, S. Ceccacci, M. Chapelle, S. Pierre-Olivier, V. Demichev, I.C. Guerrera, Neat plasma proteomics: getting the best out of the worst, Clin Proteomics 21(1) (2024) 22.

[31] P.E. Geyer, E. Voytik, P.V. Treit, S. Doll, A. Kleinhempel, L. Niu, J.B. Muller, M.L. Buchholtz, J.M. Bader, D. Teupser, L.M. Holdt, M. Mann, Plasma Proteome Profiling to detect and avoid sample-related biases in biomarker studies, EMBO Mol Med 11(11) (2019) e10427.

[32] J.F. Krey, P.A. Wilmarth, J.B. Shin, J. Klimek, N.E. Sherman, E.D. Jeffery, D. Choi, L.L. David, P.G. Barr-Gillespie, Accurate label-free protein quantitation with high- and low-resolution mass spectrometers, J Proteome Res 13(2) (2014) 1034–1044.

[33] J.R. Wisniewski, H. Koepsell, A. Gizak, D. Rakus, Absolute protein quantification allows differentiation of cell-specific metabolic routes and functions, Proteomics 15(7) (2015) 1316–25.

[34] J.R. Wisniewski, M.Y. Hein, J. Cox, M. Mann, A “proteomic ruler” for protein copy number and concentration estimation without spike-in standards, Mol Cell Proteomics 13(12) (2014) 3497–506.

[35] A. Bairoch, The Cellosaurus, a Cell-Line Knowledge Resource, J Biomol Tech 29(2) (2018) 25–38.

[36] F. Petralia, N. Tignor, B. Reva, M. Koptyra, S. Chowdhury, D. Rykunov, A. Krek, W. Ma, Y. Zhu, J. Ji, A. Calinawan, J.R. Whiteaker, A. Colaprico, V. Stathias, T. Omelchenko, X. Song, P. Raman, Y. Guo, M.A. Brown, R.G. Ivey, J. Szpyt, S. Guha Thakurta, M.A. Gritsenko, K.K. Weitz, G. Lopez, S. Kalayci, Z.H. Gumus, S. Yoo, F. da Veiga Leprevost, H.Y. Chang, K. Krug, L. Katsnelson, Y. Wang, J.J. Kennedy, U.J. Voytovich, L. Zhao, K.S. Gaonkar, B.M. Ennis, B. Zhang, V. Baubet, L. Tauhid, J.V. Lilly, J.L. Mason, B. Farrow, N. Young, S. Leary, J. Moon, V.A. Petyuk, J. Nazarian, N.D. Adappa, J.N. Palmer, R.M. Lober, S. Rivero-Hinojosa, L.B. Wang, J.M. Wang, M. Broberg, R.K. Chu, R.J. Moore, M.E. Monroe, R. Zhao, R.D. Smith, J. Zhu, A.I. Robles, M. Mesri, E. Boja, T. Hiltke, H. Rodriguez, B. Zhang, E.E. Schadt, D.R. Mani, L. Ding, A. Iavarone, M. Wiznerowicz, S. Schurer, X.S. Chen, A.P. Heath, J.L. Rokita, A.I. Nesvizhskii, D. Fenyo, K.D. Rodland, T. Liu, S.P. Gygi, A.G. Paulovich, A.C. Resnick, P.B. Storm, B.R. Rood, P. Wang, N. Children’s Brain Tumor, C. Clinical Proteomic Tumor Analysis, Integrated Proteogenomic Characterization across Major Histological Types of Pediatric Brain Cancer, Cell 183(7) (2020) 1962–1985 e31.

[37] S.J. Pelletier, M. Leclercq, F. Roux-Dalvai, M.B. de Geus, S. Leslie, W. Wang, T.T. Lam, A.C. Nairn, S.E. Arnold, B.C. Carlyle, F. Precioso, A. Droit, BERNN: Enhancing classification of Liquid Chromatography Mass Spectrometry data with batch effect removal neural networks, Nat Commun 15(1) (2024) 3777.

