## Supplementary Notes for "ibaqpy: A scalable Python package for baseline quantification in proteomics leveraging SDRF metadata"

**Table of Contents**

**Supplementary Note 1: PXD042233 data analysis** ... 3

**Supplementary Note 2: Feature intensity distribution and Copy cell number dataset PXD030306** ... 6

**Supplementary Note 3: Sample count vs coefficient of variation** ... 8

**Supplementary Note 4: DuckDB integration in ibaqpy** ... 9

**Supplementary Note 5: Evaluation of batch effect correction for Hela corpus.** ... 10

**References** ... 11

**Supplementary Note 1: PXD042233 data analysis**

In 2024, Webel et al. published a substantial collection of Hela measures acquired over approximately seven years from 2013 to 2020 [1]. The original dataset (PXD042233) comprises 7,444 MS runs from multiple instruments, including Orbitrap Q Exactive, Orbitrap Q Exactive HF, and Orbitrap Fusion Lumus. The number of peptides identified in the original study varies from 15000 to 54316 (**Supplementary Figure 1 top left**). Due to the substantial diversity, with nearly fourfold differences between MS runs, we aim to reduce the number of MS runs in our reanalysis by selecting comparable samples (MS runs) based on quality control (QC) metrics. The PRIDE Archive can download the submission QC matrix (https://ftp.pride.ebi.ac.uk/pride/data/archive/2023/12/PXD042233/pride_metadata.csv).

We performed a clustering analysis of the MS runs using the following properties: *Number of MS1 spectra*, *Number of MS2 spectra*, *MS min RT*, *MS max RT*, *MS min MZ*, *MS max MZ*, *Number of scans*, *MS/MS Submitted*, *Mass Standard Deviation [ppm]*. We developed a Python script to analyze the QC metrics (https://gist.github.com/ypriverol/5185a7c7e2954ef360037b97b5d41f52 ). In summary, the tool loads all the features for each MS run and scales each feature (StandardScaler - https://scikit-learn.org/stable/modules/generated/sklearn.preprocessing.StandardScaler.html ). The deep learning component applies an autoencoder architecture to capture non-linear patterns in scaled QC metrics. The autoencoder is implemented using PyTorch, with a three-layer structure: an input layer matching the number of features, a hidden layer with 64 neurons using ReLU activation, and a bottleneck layer with 2 neurons to reduce dimensionality. After training, the decoder reconstructs the data to minimize reconstruction error. The optimizer used is Adam, with a learning rate of 0.001, and the loss function is a mean squared error (MSE). The model is trained over 50 epochs with a batch size of 32, balancing computational efficiency and convergence. After training, the 2-dimensional latent space representation is extracted for further clustering using the k-means algorithm with 3 clusters (**Supplementary Figure 1 top right**). Figure 2 shows that the 3 clusters are not related to the year of data acquisition; MS runs acquired from different years are clustering together.

We plot the distribution of peptides identified by cluster (**Supplementary Figure 1, bottom left**), illustrating that cluster 0 has the greatest number of identified peptides, while clusters 1 and 2 have, on average, fewer identified peptides (particularly cluster 2). **Supplementary Figure 1**, bottom right, presents the final distribution of peptides by MS runs, indicating that the majority of MS runs with fewer peptide identifications have been excluded.

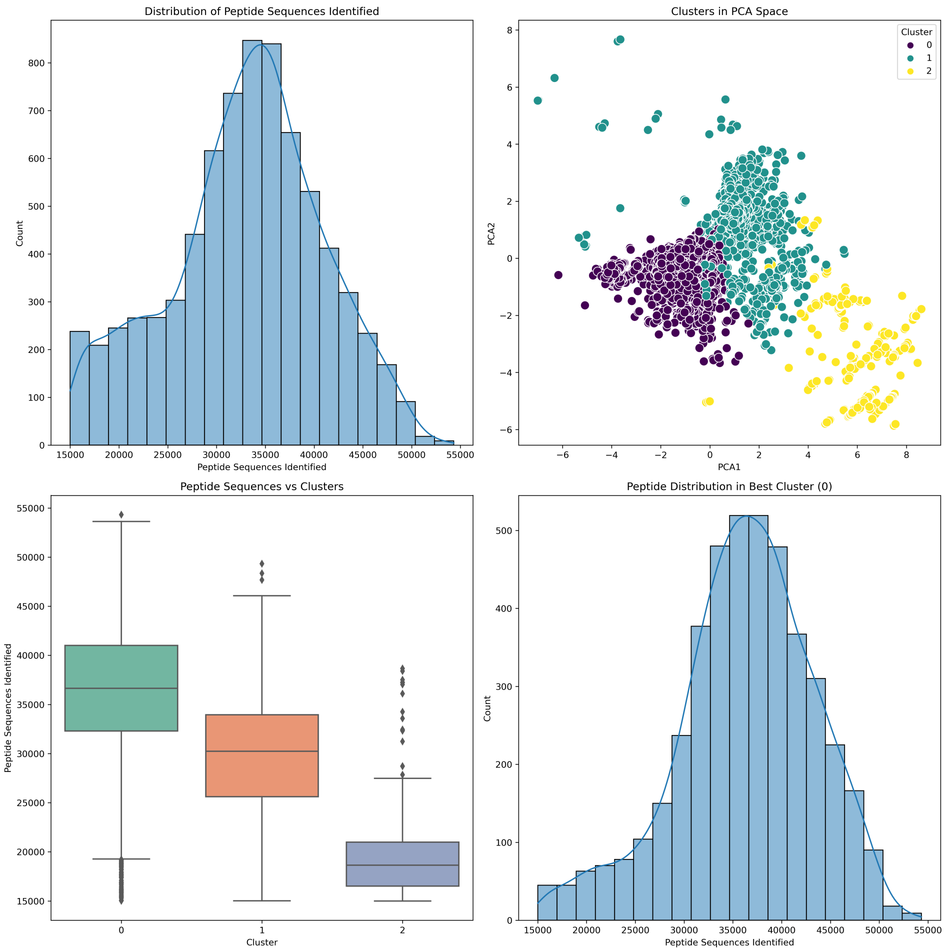


**Supplementary Figure 1**: Analysis of the MS runs QC metrics of the dataset PXD042233, and peptide sequences identified. (**top left**) The histogram represents the distribution of peptide sequences identified by MS run, showing a Gaussian-like trend from 15000 - 54316. (**top right**) The principal component analysis shows the results of the k-mean clustering (n = 3), for the k-mean MS runs were clustered using multiple properties in the QC metrics including number MS and MS/MS, minimum and maximum RT, among other properties excluding the number of peptide sequence identified. (**bottom left**) Boxplot of peptide sequences identified across clusters demonstrates differences in distribution, with Cluster 0 having the highest median value. (**bottom right**) A focused histogram of peptide sequences in the best-performing cluster (Cluster 0) illustrates its distribution.

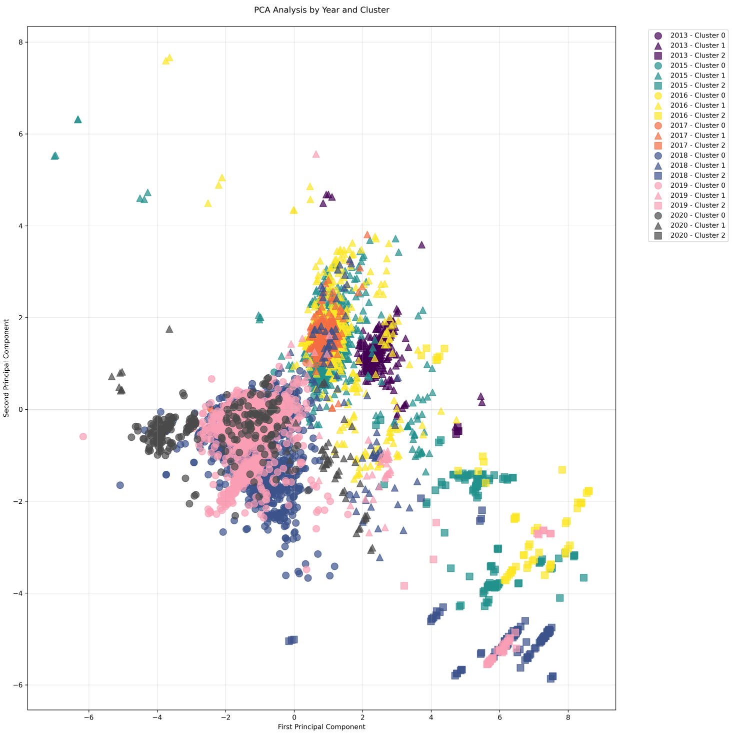


**Supplementary Figure 2**: Principal component showing the results of the cluster tag by years. The MS runs were acquired from 2013 – 2020.

**Supplementary Note 2: Feature intensity distribution and Copy cell number dataset PXD030306**

**Supplementary Figure 3** presents box plots depicting the log2-transformed intensity values for 16 samples from the **PXD030306** dataset, both before and after normalization. Each box plot corresponds to a specific sample, with the number of identified features and coefficient of variation (CV). In the pre-normalization plot, variability between samples is evident, while the post-normalization plot demonstrates improved alignment across samples, indicating successful normalization of intensity values. These figures highlight the effectiveness of the normalization process in reducing technical variability and ensuring comparability across the dataset. **Supplementary Figure 4** shows the variability of Copy cell number across samples for project PXD030306.

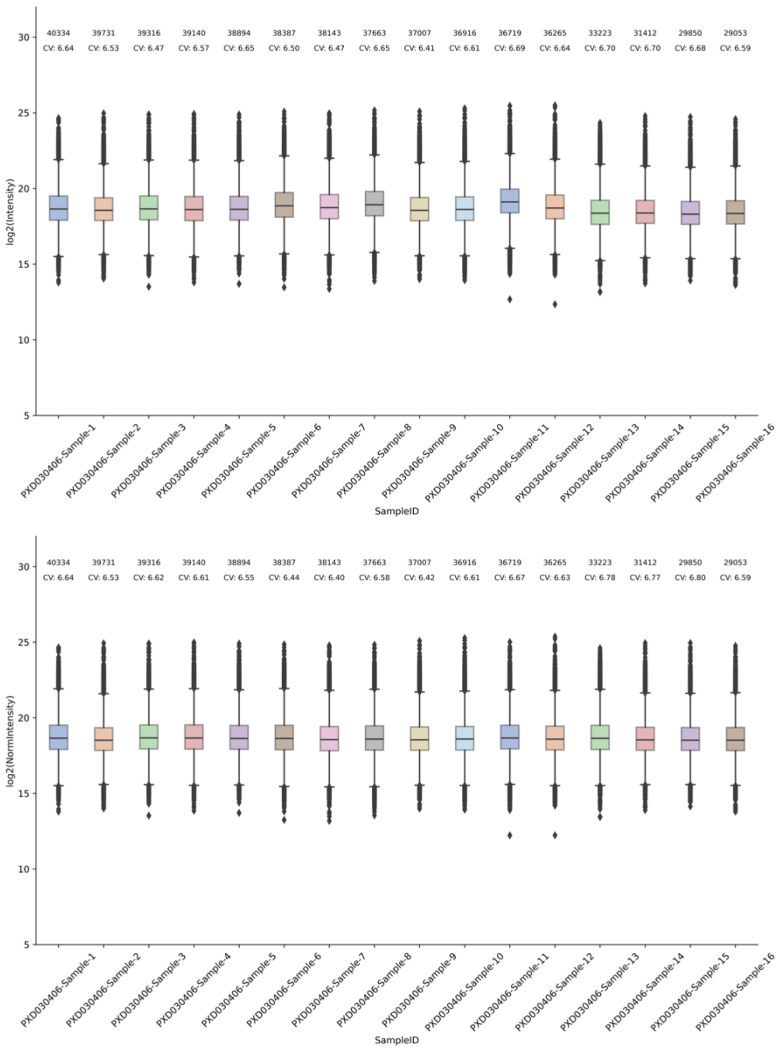


**Supplementary Figure 3**: Boxplot of the feature intensities by samples of the project PXD030306, (**top**) intensities without normalization; (**bottom**) normalized intensities.

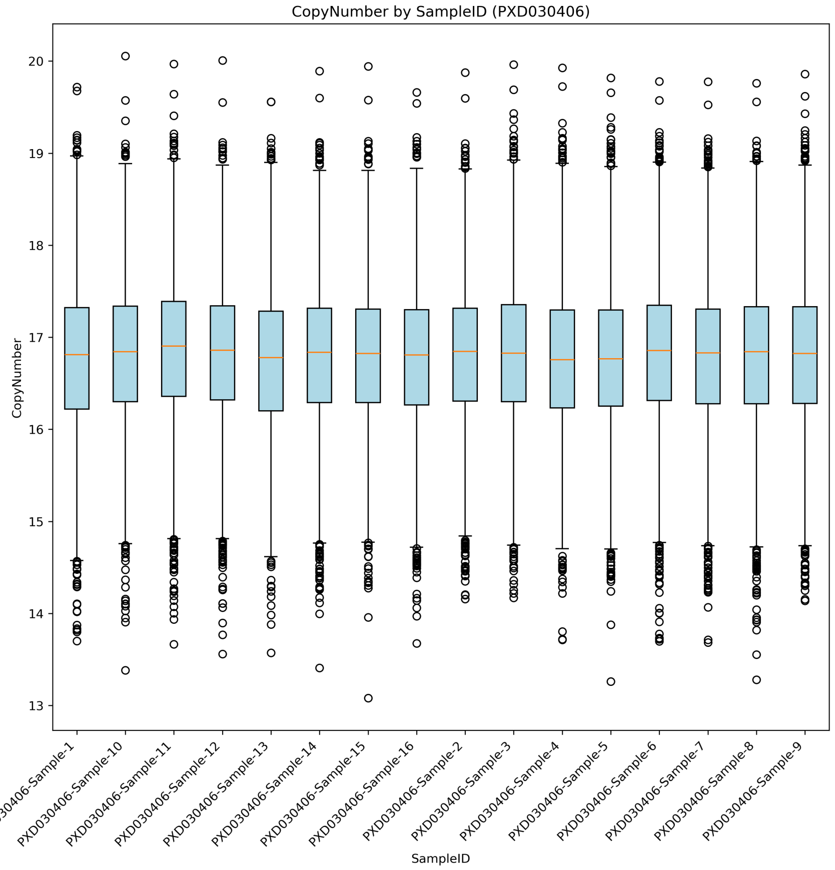


**Supplementary Figure 4**: Box plots visualise the variability of protein Cell Copy Number (Copy Number) for all samples in project PXD030406 within each group.

**Supplementary Note 3: Sample count vs coefficient of variation**

**Supplementary Figure 5** illustrates the relationship between the number of samples used in protein quantification experiments and the coefficient of variation (CV) of the quantified proteins in HeLa cells. As expected, the CV generally decreases as the sample size increases. This trend suggests that proteins quantified with higher sample numbers exhibit lower variability, indicating more reliable and consistent quantification. Conversely, proteins quantified with fewer samples tend to have higher CV, suggesting greater variability and potentially less accurate quantification. This observation aligns with the expectation that increasing the sample size can improve the precision and reliability of protein quantification, especially for proteins that are inherently more difficult to quantify due to low abundance or other factors.

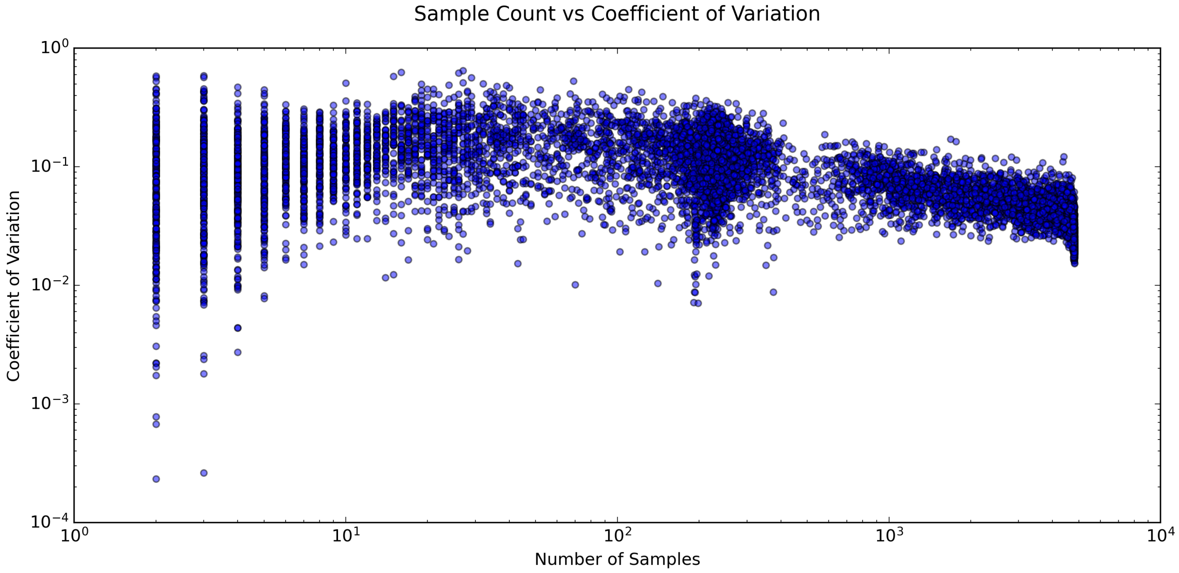


**Supplementary Figure 5**: Coefficient of variation by protein versus the number of samples quantified.

**Supplementary Note 4: DuckDB integration in ibaqpy**

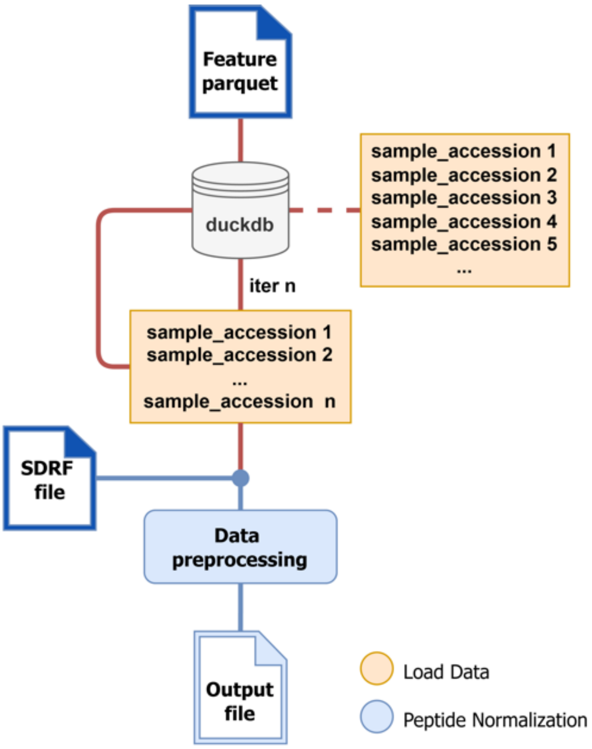


**Supplementary Figure 6**: Peptide features data is stored in a parquet-based file (https://github.com/bigbio/quantms.io) and processed in batches, and it is queried using DuckDB for efficient handling of large datasets.

### Supplementary Note 5: Evaluation of batch effect correction for Hela corpus.

**Supplementary Table 1**: Comparison of Silhouette Score and Entropy of Batch Mixing (EBM) before and after correction. A decrease in the Silhouette Score indicates successful mitigation of batch effects, while an increase in EBM suggests improved mixing of samples across batches. Both scores were computed with the function *"cohort_qc"* from *inmoose*. EBM was calculated using 10 principal components (PCs) and *k*=200 nearest neighbours. Maximum entropy is equal to log2(*B*), where *B* is the number of batches [2].

| Metric | Before Correction | After Correction | Scales* | Interpretation |
|---|---|---|---|---|
| **Silhouette Score** | -0.0643 | -0.2834 | [-1, 1] | Decrease suggests successful batch effect mitigation. |
| **Entropy of Batch Mixing** | 0.1521 | 0.4339 | [0, log2(*B*)] | Increase suggests better mixing of batches. |

### References

[1] H. Webel, Y. Perez-Riverol, A.B. Nielsen, S. Rasmussen, Mass spectrometry-based proteomics data from thousands of HeLa control samples, Sci Data 11(1) (2024) 112.
[2] A. Droit, S. Pelletier, M. Leclerq, F. Roux-Dalvai, M. de Geus, S. Leslie, W. Wang, T. Lam, A. Nairn, S. Arnold, B. Carlyle, F. Precioso, Enhancing Classification of liquid chromatography mass spectrometry data with Batch Effect Removal Neural Networks (BERNN), Res Sq (2023).
